# Vitamin B12 partially rescues embryonic cell migration defects in *C. elegans* ephrin mutants by improving both propionic acid breakdown and one-carbon cycle metabolic pathways

**DOI:** 10.1101/2024.11.05.622144

**Authors:** Tushar H. Ganjawala, Erin Hsiao, Prativa Amom, Radmehr Molaei, Samantha Goodwin, Amanda L. Zacharias

## Abstract

Successful cell migration followed by tissue fusion is required for organogenesis in a number of tissues, many of which are susceptible to gene-environment interactions that can result in congenital anomalies. In *C. elegans* embryogenesis, one such event is the closure of the ventral cleft, which must occur as the first step in morphogenesis, otherwise embryos arrest; this process depends on ephrin signaling, but no single gene mutation is fully penetrant embryonic lethal. We exposed hermaphrodites mutant for *vab-1*, the *C. elegans* ephrin receptor, to various environmental conditions and found that vitamin B12 supplementation could partially rescue the embryonic lethality of multiple alleles from 40% survival to 63% survival. Analysis of *vab-1* mutant phenotypes showed that vitamin B12 reduced the frequency of ventral cleft closure failure by promoting more normal cell positions and increased migration by the precursor cells of those that adhere to close the cleft. We found that vitamin B12 partially rescued the embryonic lethality of other ephrin pathway mutants, but not mutants that have ventral cleft defects due to cell adhesion or cell fate defects. We found that rescue by vitamin B12 depends on its functions in both mitochondrial propionic acid breakdown and the one-carbon cycle, and that antioxidant treatment can also partially rescue ephrin pathway mutants. These results are distinct from the larval response to vitamin B12, which depends only on the one-carbon cycle, emphasizing the unique metabolism of embryos and particularly the metabolic needs of migrating cells. Overall, our findings highlight the *C. elegans* embryo as a model system to investigate gene-environment interactions and developmental metabolism.

## INTRODUCTION

Cell migration followed by tissue fusion is critical for the development of many organisms and plays a critical role in organogenesis. In this process, two layers of migrating cells come together, often at the midline, and adhere to each other to form a newly unified layer of cells. In the embryonic development of the nematode worm, *C. elegans*, the first tissue fusion event that occurs is the closure of the ventral gastrulation cleft: as gastrulating pharyngeal and muscle precursor cells internalize, they leave a cleft on the ventral side of the embryo that neuronal and epithelial precursors from the left and right edges fill by migrating toward the midline and adhere to each other^1^. This process occurs between the 190 cell stage and 350 cell stage (170-250 Sulston minutes) and includes one round of cell divisions by the migrating cells, although they maintain the anterior-posterior division orientation induced by Wnt signaling^2–4^. The newly formed epithelial layer later serves as a substrate for the migration of the skin cells during ventral enclosure (at ∼300 Sulston minutes); if ventral cleft closure fails, ventral enclosure subsequently fails and the embryos burst open during morphogenesis between comma and 1.5 fold stage and fail to hatch^1^.

Numerous signaling pathways have been identified that contribute to the closure of the cleft. A key signaling pathway required is eph-ephrin signaling, although there is additional redundancy with plexin-semaphorin signaling: embryos mutant for the sole ephrin receptor, *vab-1*, fail to hatch 60% of the time, while double mutants with the plexin, *mab-20*, are fully penetrant embryonic lethal^5–7^. The Robo receptor, *sax-3*, also contributes, but it may be activated by binding to the atypical ephrin, *eph-4*, as recent biochemical evidence shows the two can interact and its canonical slit ligand, *slt-1*, is not expressed or required^7–11^. Double mutants for *vab-1* and *sax-3* are also fully penetrant embryonic lethal^9^. For all of these signaling pathways, none of the loss-of-function mutants are fully penetrant lethal on their own, indicating that there is sufficient redundancy to promote cleft closure when one pathway is disrupted. We recently found that appropriate cell fate specification of the neuroblasts, muscle, intestinal and hypodermal cells are all separately required for normal ventral cleft closure (Ganjawala, Amom, et al., in preparation), although the extent to which these disruptions impact cell signaling is unclear. However, it emphasizes the complexity of this process.

In addition to the signals that presumably provide direction for the migration of the cells that close the cleft, these must have sufficient energy and nutrients to carry out this migration. During *C. elegans* embryogenesis, all energy and nutrients must come from what was maternally deposited in the egg prior to fertilization, as the embryo cannot eat until it hatches. Recently, it has been found that the laboratory diet of *E. coli*, which was selected for convenience and is not a food source that *C. elegans* encounters in its natural environment, is depleted in the nutrient vitamin B12 (cobalamin). Vitamin B12 is a necessary co-factor for two key metabolic processes, breakdown of propionic acid in the mitochondria, which also generates succinyl-CoA that can be used to produced additional energy, and the one carbon cycle, which produces methyl donors for modifying proteins, nucleic acids, and lipids^12^. In a series of seminal papers, the Walhout group found that *C. elegans* larva supplemented with vitamin B12, either in the form of a chemical additive or bacteria that naturally produce it, passage through the larval stages is accelerated because of the increased availability of methyl donors^13–15^. On a diet low in vitamin B12 like OP50 *E. coli, C. elegans* larva can avoid the buildup of propionic acid by activating an alternative shunt pathway to break it down that produces ketone bodies as an energy source^16–18^. Vitamin B12 is a compound produced only by some bacterial species^12,15^, so *C. elegans* likely evolved this shunt to be able to colonize rotting vegetation with low fractions of vitamin B12-producing bacteria. The impact of vitamin B12 on *C. elegans* embryonic development has been largely unexplored.

Human cellular metabolism likewise requires vitamin B12 as a co-factor for critical metabolic reactions in the one carbon cycle and propionic acid breakdown, and evidence indicates patients with inborn errors of metabolism utilize a homologous propionic shunt pathway^19^. Evidence from human embryonic development indicates that vitamin B12 is required for several key developmental events. Maternal deficiency in vitamin B12 has been linked to defects in neural tube closure, a tissue fusion event, and heart remodeling, which involves cell migration and tissue fusion^20^. Another micronutrient that contributes to the one-carbon cycle, folate, is also implicated in neural tube closure defects, but not heart defects^20^, which suggests that cobalamin’s primary function in neural tube development may be in the one carbon cycle, while its primary function in heart development may be in mitochondrial propionic acid breakdown, although this remains to be determined empirically.

Here, we investigate the ability of different environmental exposures to impact ventral cleft closure in *vab-1* mutant embryos and find that vitamin B12 treatment of hermaphrodites has a significant ability to increase survival of their offspring. Vitamin B12 can rescue mutants in the Ephrin, Robo, and Plexin signaling pathways. We find that this vitamin B12 rescue requires the function of both the one-carbon cycle and propionic acid breakdown pathways. Consistent with this, we find that folate supplementation or antioxidant treatment also result in partial rescue. This indicates that the essential role for vitamin B12 in cellular metabolism is distinct between embryos and larva, and emphasizes the distinct metabolic needs of migrating cells in embryonic development.

## RESULTS

To determine the impact of different environmental conditions on ventral cleft closure, we evaluated their impact on the hatch rate of *vab-1* mutant embryos, which encodes the sole receptor for Ephrin, a signaling pathway redundantly required for this embryonic tissue fusion event. In comparison to elevated glucose exposure, elevated temperature and ethanol exposure, we found that vitamin B12 supplementation of L4 worms significantly and consistently increased the hatch rate of their subsequent offspring (referred to hereafter as parental supplementation) (Figure 1A). Approximately 40% of *vab-1(e2027)* loss of function mutant embryos hatch when worms are grown on normal growth media (NGM), consistent with previous reports^6^. We found the hatch rate increased to 63% with a parental vitamin B12 supplement. Although the embryonic lethality of these mutants was not completely rescued, this represents a significant population effect for the mutant worms: due to exponential growth a mutant population that had B12 supplementation would outnumber one grown on NGM 100 to 1 after just 11 generations. We found a similar effect by switching the bacterial diet from OP50 *E. coli* to a *Comamonas* strain, DA1877, which naturally produces vitamin B12^15^ (Figure 1A). We also observed significant but partial rescue for other loss of function and intermediate function mutations in *vab-1*^5^ (Figure 1B). The impact of vitamin B12 on *vab-1* mutant embryo survival showed a dose-response curve (Figure 1C), but it did not rescue the neuronal pathfinding defects reported in larva or the defects in skin development that result in abnormal head morphology at the standard dose (Figure S1). To determine if the effect on embryonic survival could be due to accelerated parental development caused the vitamin B12 supplement, we used serial transfer to determine the hatch rate of mutant hermaphrodites for the first three days after L4 stage and found the same rescue effect on each day (Figure 1D). This indicates that the impact of vitamin B12 is on the embryos, not the relative age of the parent.

**Figure 1.**
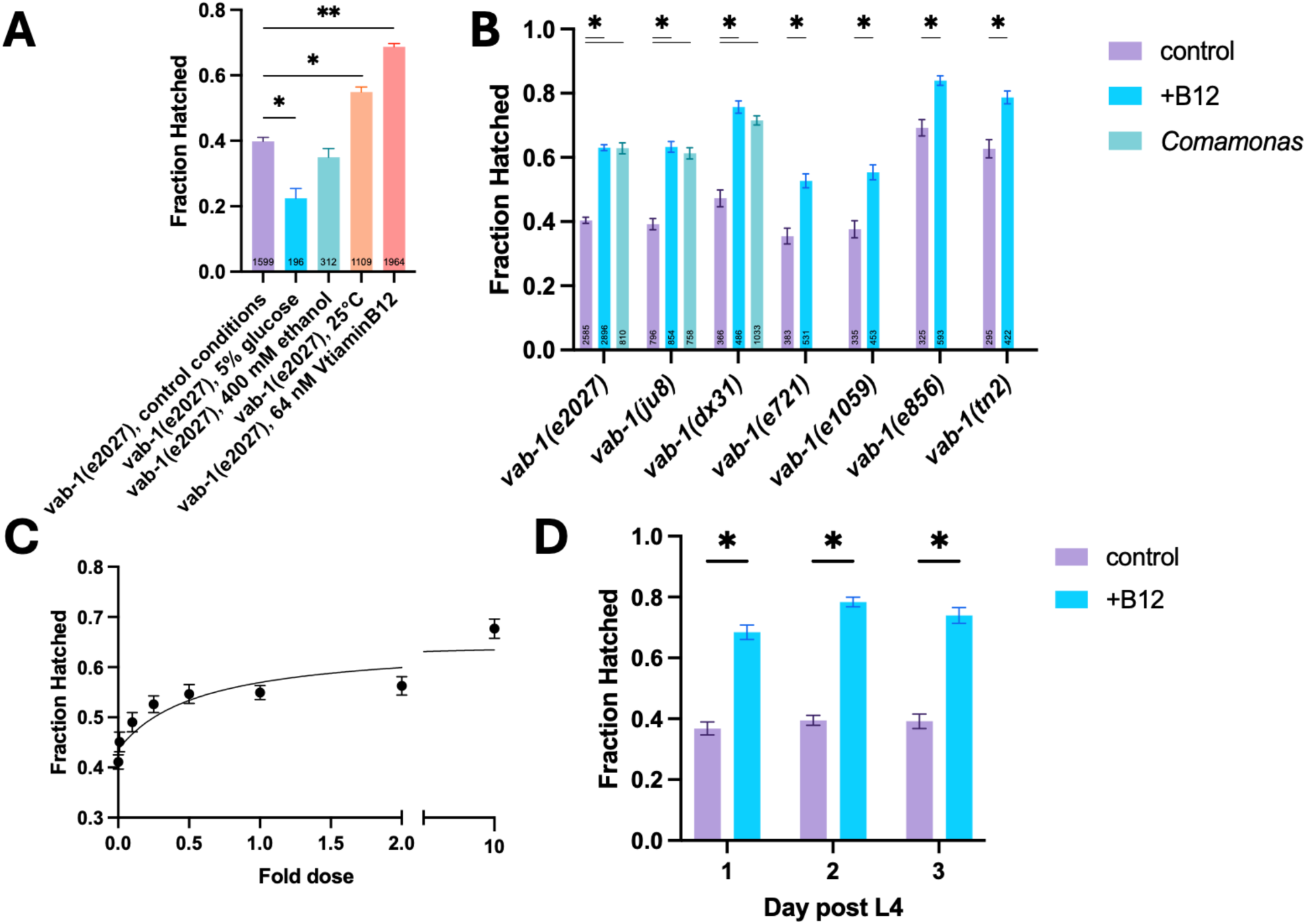
Vitamin B12 partially rescues embryonic lethality of *vab-1* ephrin receptor mutant embryos. A) While other environmental perturbations affected the fraction of *vab-1(e2027)* embryos that hatched, parental vitamin B12 supplementation showed a robust rescuing effect that prompted further investigation. B) Vitamin B12 supplementation or *Comamonas* bacteria, which naturally produce vitamin B12, partially rescue different null (*e2027, ju8, dx31, e721, e1059*) and partial loss of function (*e856, tn2*) alleles of *vab-1*. C) The fraction of *vab-1(e2027)* embryos hatched increases with the concentration of vitamin B12 supplemented in a dose-dependent manner. D) Rescue of *vab-1(e2027)* mutant embryos by vitamin B12 does not depend on how recently the parent started laying eggs.

To better understand the role of VAB-1-mediated signaling on ventral cleft closure, and how this might be impacted by the addition of vitamin B12, we first utilized several embryonic single-cell RNA-seq datasets to clarify which cells were competent to send and receive ephrin signals^11,21,22^. While mRNA expression is not directly indicative of protein expression, we find that *vab-1, vab-2, efn-4, mab-20, ptp-3*, and *sax-3* are maternally deposited, later degraded, and dynamically regulated throughout embryogenesis (Supplemental Table 1). Ligand *efn-2* is also dynamically and specifically expressed, while *efn-3* is not observed. By the time ventral cleft closure begins, almost all cells are expressing ligand, receptor, or both, for both the VAB-1/VAB-2/EFN-2 and SAX-3/EFN-4 combinations (Figure 2A, Supplemental Table 1). This suggests that rather than providing strong directionality to cell migration, the purpose of these signals may be to trigger migratory behavior.

**Figure 2.**
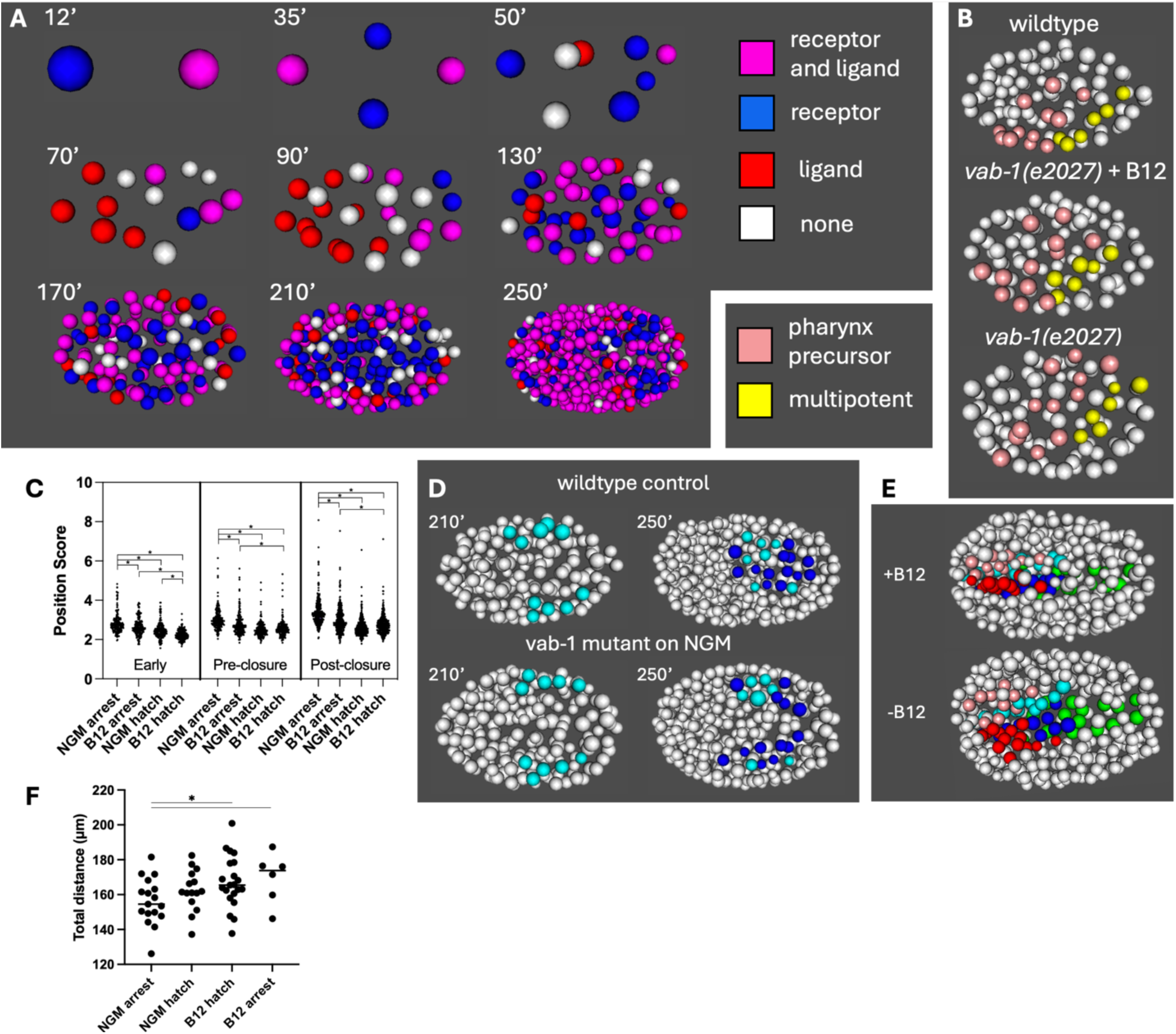
Vitamin B12 rescues *vab-1* mutant embryos by improving cell positions and migration of the ventral cleft cells. A) Expression of ephrin receptor and ligand mRNA is widespread and dynamic throughout embryogenesis. Here the ephrin receptor is *vab-1* and ligand is *vab-1* or *efn-2*. B) *vab-1(e2027)* mutant embryos show displacement of certain pharynx (pink) and multipotent (grey) precursor cells, but displacement is less severe in vitamin B12 treated embryos. C) Position scores are worse for the cells of arresting embryos in the early phase of embryogenesis and worsen as development progresses, regardless of B12 treatment. Normal Growth Media (NGM) indicates standard food. The position score metric reflects the displacement of the nucleus from its normal position combined with the level of disrupted neighbor relationships. Each point represents one cell from one embryo (n_embryos_ = 17, 6, 15, 21) D) Prior to cleft closure, precursor cells (light blue) adopt normal positions in *vab-1(e2027)* mutant embryos but the daughters of these cells fail to close the cleft fourty minutes later. Dark blue cells are among the top 5% most disrupted cell positions. E) A B12 supplemented *vab-1(e2027)* embryo shows normal pharyngeal organization, with tightly ordered nuclei on the left (AB-derived: pink, MS-derived: light blue) and right (AB-derived: red, MS-derived: blue), while the unsupplemented embryo shows displacement and disorganization. E) The cleft precursor cells of B12 treated cells show a longer migration path. Each point represents the summed path for the cleft precursor cells of one embryo (n_embryos_ = 17, 15, 21, 6).

To further determine the role of vitamin B12 in improving the viability of *vab-1* mutant embryos, we used a fluorescent live imaging approach to track nuclei positions across embryonic development, through the 350 cell stage when ventral cleft closure has completed and compared them to a wildtype model^4,23,24^. Distinct from its role in larval development, we did not observe an increase in developmental rate as measured by the length of each cell cycle in either wildtype or *vab-1* mutant embryos supplemented with B12 (Supplemental Tables 2 and 3). Furthermore, we found that vitamin B12 reduced the fraction of embryos that had ventral cleft closure defects (17/32 on standard conditions vs. 6/27 with vitamin B12 supplement.

Consistent with a potential early role for ephrin signaling, we identified cell positions that were significantly disrupted in early development of *vab-1* mutant embryos as early as the 26 cell stage (Figure 2B). These include a number of MS-derived cells as well as AB-derived cells fated to generate pharyngeal cells (Supplemental Table 4). Some of these cells have more normal positions in embryos that hatch, particularly in those supplemented with vitamin B12. We developed a cell “position score” metric which is the sum of the nuclear displacement from the wildtype position in microns and the cell’s nearest neighbor score, in which higher values indicate greater distance from its typical neighboring nuclei^23,24^. For all embryos analyzed, cells of arresting embryos on a control diet have the worst scores, while those of arresting B12 supplemented embryos also score poorly (Figure 2C). In contrast, cells of hatching B12 supplemented embryos have the lowest position scores at all timepoints. The cells of embryos on a control diet that do go on to hatch have significantly better position scores early on (<100 cells), and they are similar to the hatching B12 supplemented embryos at later timepoints. This indicates that more severe defects in early developmental stages are more likely to result in failure to hatch. We also directly compared control embryos to vitamin B12 embryos to identify cells for which the position score experienced the greatest rescue by vitamin B12 supplementation; these also included a number of pharyngeal precursors (Supplemental Table 5). By the end of the period in which the ventral cleft normally closes (350 cell stage, 250’), we observe that the cells that normally close the cleft are among those most displaced from their normal position (Figure 2D). Other severely displaced cells are future pharyngeal cells, and pharyngeal organization is severely disrupted in un-supplemented embryos that go on to arrest (Figure 2E). To determine if vitamin B12 was playing a role in promoting migration of the cells that close the cleft, we conducted a non-directional path length analysis on the inner cleft precursor cells that undergo the longest migration (Supplemental Table 6). We find that vitamin B12 promotes a longer migration path for these cells regardless of whether the embryos go on to hatch (Figure 2F). Thus, we conclude that parental vitamin B12 supplementation is at least able to partially rescue some early cell position defects and promote a longer migration for cells that close the ventral cleft.

To determine if other ventral cleft closure mutants were impacted by vitamin B12 supplementation, we first examined ephrin ligand mutants. *C. elegans* has four ephrin ligand genes, *vab-2* (AKA *efn-1), efn-2, efn-3, efn-4*, although *efn-3* is not expressed during embryogenesis. The ligand mutants have varying degrees of embryonic lethality, never more than the receptor itself, and all were significantly but not completely rescued by vitamin B12 (Figure 3A). We also examined mutants from other signaling pathways that contribute to ventral cleft closure, and we found that vitamin B12 supplementation could also significantly but incompletely rescue the embryonic lethality of *sax-3/Robo* and *mab-20*/*Plexin* (Figure 3A). We also used vitamin B12 to supplement balanced worms carrying a weak allele of *vab-1* and a loss of function allele of *sax-3*, but we never observed any hatched L1s that lacked the marked balancer, indicating that vitamin B12 is unable to rescue a completely penetrant lethal combination of signaling mutations. We used vitamin B12 to supplement mothers of other strains with partial embryonic lethality due in part to defects in ventral cleft closure including *hmp-1/a-catenin*, which is required for cell adhesion after closure, and *hlh-1/MyoD1* and a triple mutant for *cnd-1/NeuroD; ngn-1/Neurog; lin-32/Atoh*, which are required for cell fates that promote ventral cleft closure (Ganjawala, Amom, et al., in preparation), but observed no change in the rate of embryonic lethality (Figure 3B).

**Figure 3.**
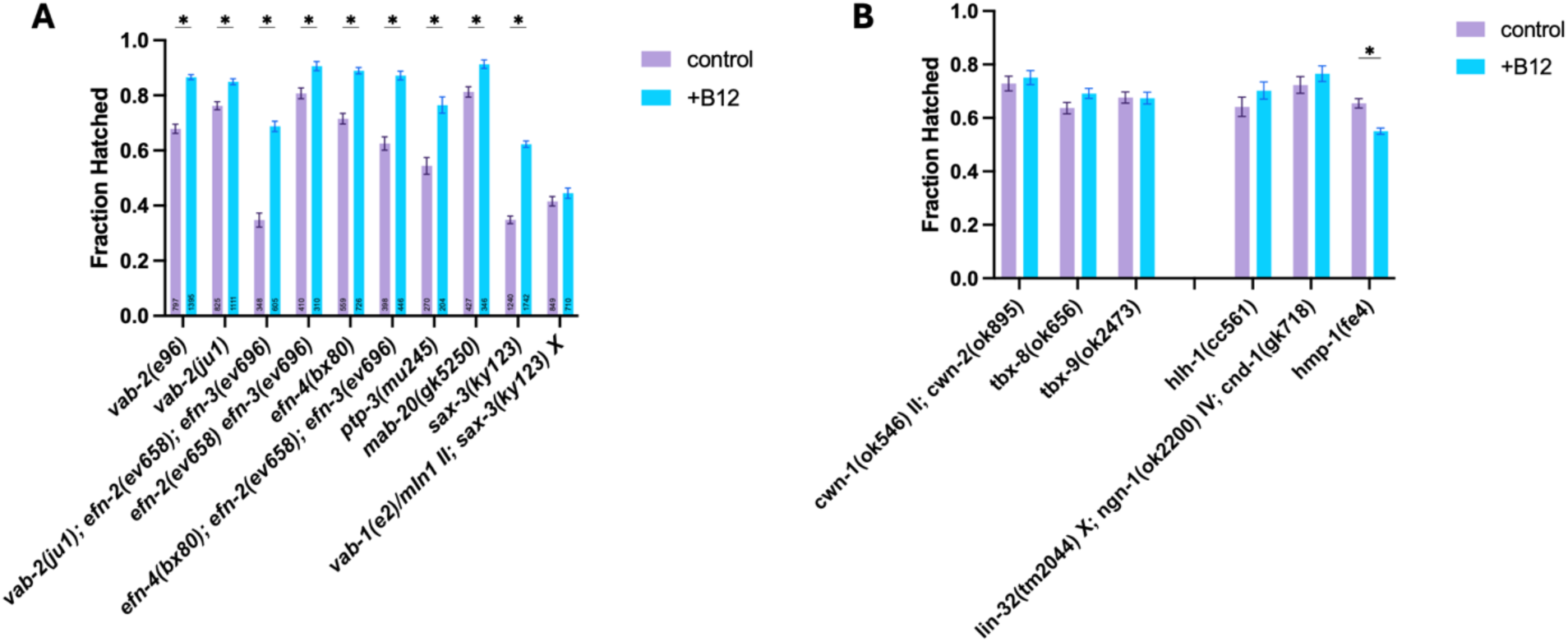
Vitamin B12 can rescue Ephrin ligand and other related mutants but not embryos that arrest due to other causes. A) Vitamin B12 partially rescues ephrin ligand mutants (*vab-2, efn-2, efn-3, efn-4)*, an associated phosphatase *ptp-3*, semaphorin *mab-20*, and robo receptor *sax-3*, but cannot rescue a balanced lethal *vab-1; sax-3* double mutant strain. B) Vitamin B12 does not rescue embryonic lethality in Wnt ligand *cwn-1;cwn-2* or *tbx-8* or *tbx-9* transcription factor mutants. Vitamin B12 also does not rescue the partial embryonic lethality of other mutants shown to have partially penetrant ventral cleft closure defects due to cell fate abnormalities (*hlh-1* and *lin-32;ngn-1;cnd-1*) or loss of cell adhesion (*hmp-1)*.

Furthermore, we saw that parental vitamin B12 supplementation did not alter the rate of embryonic lethality due to mutations in other unrelated genes, such as the transcription factors *tbx-8* and *tbx-9*, or two Wnt ligands expressed in mid-embryogenesis, *cwn-1; cwn-2* (Figure 3B). This indicates that the rescuing effect of vitamin B12 is rather limited to defects in ventral cleft closure caused by disruptions in signaling and it can never completely rescue embryonic lethality, even in these cases.

To further confirm the reduction in embryonic lethality required vitamin B12, we evaluated whether disruption of vitamin B12 transport or processing could prevent vitamin B12 rescue of *vab-1* mutants. To do this, we generated worms double mutant for *vab-1* and the vitamin B12 processing enzyme *cbcl-1*, which converts cobalamin (B12) to Ado-cobalamin, the bio-available form we generally use for supplementation. These worms were supplemented with cobalamin and we observed no rescue (Figure 4A). Similarly, we used RNAi to knock-down *mmad-1*, previously known as Y76A2B.5, which is required for vitamin B12 processing and transport, in *vab-1* mutant worms and found it was also required for embryonic rescue relative to RNAi controls (Supplemental Figure S4A). Since this gene is the homolog of the human gene MMADHC, which is associated with methylmalonic aciduria and homocysturia, cblD type, we named this gene *mmad-1*.

**Figure 4.**
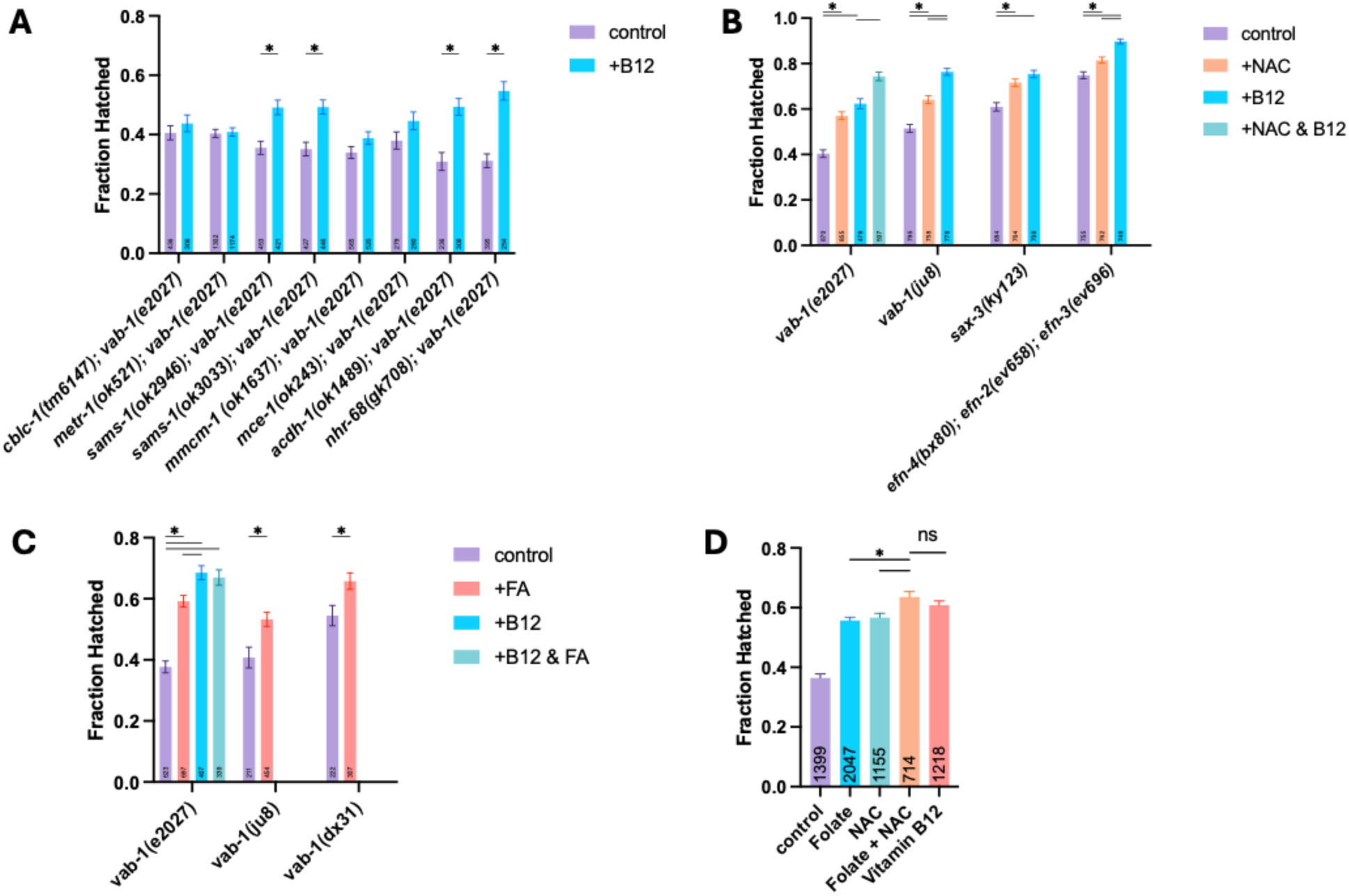
Rescue of *vab-1* ephrin mutants depends on the function of vitamin B12 in both the one-carbon cycle and propionic acid breakdown pathways. A) Loss of function mutations in some metabolic genes prevents vitamin B12 rescue of *vab-1(e2027)* mutant embryos. B) Parental treatment with the antioxidant N-acetylcysteine (NAC) can partially rescue ephrin pathway mutant embryos but not as well as vitamin B12. C) Parental supplementation with folic acid (FA) can partially rescue multiple alleles of *vab-1*, but not as well as vitamin B12. D) Parental treatment with both folate and NAC rescues an equivalent amount to vitamin B12.

To determine whether the reduction in embryonic lethality depended on the role of vitamin B12 in the one-carbon cycle or in propionic acid breakdown, we used similar double mutant and RNAi approaches. For the one-carbon cycle, we observed that although *sams-1;vab-1* double mutant embryos were able to be rescued by vitamin B12, *metr-1; vab-1* embryos were not. Further analysis showed that other homologs, *sams-3, sams-4*, and *sams-5*, are expressed during embryogenesis (Supplemental Table 7)^22^, so these genes may have redundant function. Therefore, we conclude that a functional one-carbon pathway is required for vitamin B12 rescue of *vab-1* mutant embryos. Similarly, for the propionic acid breakdown pathway, we observed that *mmcm-1; vab-1* and *mce-1; vab-1* double mutant embryos could not be rescued by vitamin B12, indicating that the primary propionic breakdown pathway is also required for rescue. A key shunt pathway member *acdh-1* and its upstream regulator *nhr-68*^14,25^ are not required for rescue by vitamin B12, in part because they are downregulated when vitamin B12 is present. Together, these results indicate that vitamin B12 improves the viability of *vab-1* mutant embryos due to its actions in both the one-carbon cycle and propionic acid breakdown metabolic pathways.

If vitamin B12 is improving propionic acid breakdown, this may improve mitochondrial function and reduce redox stress. To determine whether reduced redox stress may explain why vitamin B12 improves *vab-1* mutant embryo viability, we treated embryos with a strong antioxidant, N-acetylcysteine (NAC)(Figure 4B). We found that NAC was also able to partially rescue multiple alleles of *vab-1*, as well as *sax-3*, and *efn-4; efn-2; efn-3* mutants, but rescue was never as strong as with vitamin B12 itself, consistent with an additional role for vitamin B12. We also saw even greater levels of rescue (74% eggs hatched) when we combined NAC with vitamin B12, consistent with our previous finding that our standard dose of vitamin B12 was not achieving maximal effect. Besides removing a toxic byproduct from mitochondria, the propionic acid breakdown pathway also produces Succinyl-CoA, which produces energy through the citric acid cycle, while the shunt pathway produces energy in the form of Acetyl-CoA and ketone bodies. Recently, it was shown that the enzymes to generate ketone bodies or exogenous supplementation are required for embryogenesis in the absence of vitamin B12^18^, so we investigated whether ketone body supplementation could cause rescue of *vab-1* mutant embryos by providing an additional source of energy. We observed a mild rescue with acetoacetate, but not 3-hydroxybutyrate, and no additional rescue when combined with vitamin B12 or NAC. This indicates that if increased energy from propionic acid breakdown is involved in the rescue of *vab-1* mutant embryos by vitamin B12, ketone bodies are not a sufficient replacement.

An increased level of vitamin B12 would also permit increased flux through the one carbon cycle, because it is a critical factor for processing folate which generates purines for DNA synthesis and methyl donors for protein and DNA modification, all of which are critical for embryogenesis. We hypothesized that introducing additional folate to the system in the form of folic acid (FA) might have a similar effect. Indeed, we did see a similar partial rescue of several *vab-1* alleles with FA supplementation, but again it was not as high as vitamin B12 itself. Adding FA to vitamin B12 supplementation did not generate an additional increase in rescue, indicating that folate is not a limiting reagent when vitamin B12 is present in sufficient amounts. Given that folate and the antioxidant NAC might act separately on both pathways that vitamin B12 is critical for, we hypothesized that combining the two might achieve a similar level of rescue as vitamin B12. We observed that separately, FA and NAC each partially rescue to a similar level, and that when combined, they achieve a level of rescue indistinguishable from vitamin B12, further emphasizing that rescue depends on both pathways for which vitamin B12 is required.

## DISCUSSION

For tissue fusion events to happen appropriately during embryonic development, cells must migrate towards each other and adhere to each other. Here we examined a series of mutants in the ephrin and related signaling pathways, in which this process fails to occur some of the time due to reduced signaling activity, resulting in embryonic lethality. Since ephrin ligands and receptors are broadly expressed in the *C. elegans* embryo, fusion can happen when signaling activity is reduced due to pathway mutations, and the cells do not migrate long distances, we hypothesize that the role of ephrin signaling is primarily to trigger cell migration activity rather than to provide a directionality to migration. Here, we showed that vitamin B12 can significantly improve the success of the ventral cleft closure process during embryogenesis, resulting in increased viability of ephrin pathway embryos with genetic signaling deficits.

During embryonic development, cells undergo changes very rapidly, including rapid cell division (every ∼40 minutes in the *C. elegans* embryo), chromatin modifications during differentiation, and for some cells, migration to the appropriate location. Vitamin B12 is an important enzymatic co-factor for two metabolic processes that contribute to these events, so it is critical for normal development. Vitamin B12 is part of the one-carbon cycle which generates nucleotides for DNA synthesis and methyl donors, which are important for chromatin modifications. Since the rate of cell divisions does not change in response to vitamin B12, we hypothesize that the primary role of vitamin B12 in promoting successful ventral cleft closure is in generating methyl donors that are critical for remodeling chromatin during differentiation. Indeed, classic and more modern approaches have shown that chromatin changes related to differentiation begin at the 50-100 cell stage prior to when ventral cleft closure occurs^26–30^. We hypothesize that faster or more complete differentiation of embryonic cells promotes greater expression of the other ephrin pathway components that provide redundancy in the pathway, improving signaling and thus cell migration, which ultimately leads to increased likelihood of successful ventral cleft closure. Fitting with this, other experiments in our laboratory have found that disrupting key transcription factors that promote cell fate such as *hlh-1/MyoD* also disrupt ventral cleft closure (Ganjawala, Amom et al., in preparation). It may also explain why vitamin B12 does not rescue later larval defects in neuronal migration and head morphogenesis, as most cells are fully differentiated by this period.

Vitamin B12 is also part of the propionic acid breakdown pathway in mitochondria, which maintains mitochondrial health, reduces redox stress, and generates energy, which could help migrating cells as they are assumed to utilize more energy than those that do not migrate. If migrating cells are using more energy, they also experience more mitochondrial redox stress if propionic acid breakdown does not occur appropriately.

Consistent with this, we find that antioxidant treatment with NAC can partially rescue ephrin signaling mutant viability, suggesting that reducing mitochondrial stress may improve ventral cleft closure. Existing gene expression datasets indicate that some of the necessary genes of the propionate shunt are not robustly expressed during the early phases of embryonic development when ventral cleft closure occurs (Supplemental Table 8)^22,31,32^. This indicates that even if the propionate shunt is active, it results in suboptimal mitochondrial function. Furthermore, we also found that vitamin B12 treatment promotes increased migration distance by the cells that close the cleft, regardless of whether it successfully closes and the embryo ultimately hatches. This indicates that vitamin B12 improves the health of cells to enable them to migrate more even if they do not migrate to the appropriate positions to make the necessary cell-cell adhesions to close the cleft.

The fact that both functions of vitamin B12 are required for rescue of ephrin and related pathway mutants underscores the complexity of embryonic development. Cell health and robust signaling are both required for complex events depending on cell migration like ventral cleft closure. This fits with data from human congenital anomalies showing that both genetic and environmental factors contribute^33^. Furthermore, it highlights the differences between larval development and embryonic development. Embryonic development is not accelerated by vitamin B12, suggesting that it may already be proceeding at a maximal rate or that nucleotides, methyl donors, and energy are not rate-limiting, but provided in sufficient amounts from the oocyte. Furthermore, there are significantly fewer cell migration events during larval development and they are not required for viability, which may explain why the vitamin B12-dependent propionate breakdown pathway is not required for accelerated larval development^15,34^. Overall, our results show that the *C. elegans* embryo is a viable model system for investigating gene-environment interactions related to development and that the the low levels of vitamin B12 in standard growth conditions impacts the development of some types of mutant embryos.

## MATERIALS AND METHODS

### Worm strains and maintenance

A list of strains used can be found in Supplemental Table 9. Some strains were provided by the CGC, which is funded by NIH Office of Research Infrastructure Programs (P40 OD010440). Worms were maintained on standard NGM media with OP50 food source at 20°C unless otherwise noted. *Comamonas* DA1877 bacteria was used for some experiments on NGM plates supplemented with streptomycin. New strains were generated through standard genetic crosses and verified with PCR genotyping or phenotype analysis.

### Supplementation and drug treatment

Vitamin B12 plates: Ado-Cobalamin (Sigma C0884) was added at 64 nM final concentration(1X) unless otherwise noted to NGM agar media after cooling to 55°C. After plates were poured they were protected from light while they dried, seeded with OP50 bacteria and stored away from light. For experiments with *cblc-1;vab-1* double mutants, cyanocobalamin (Sigma V6629) was used. 5% glucose plates: Glucose solution was spread on unseeded NGM plates at a final volume of 5% by mass. Plates were dried and then seeded with OP50 for use. Ethanol plates: concentrated ethanol solution was added to unseeded NGM plates at a final concentration of 400 mM. Plates were sealed with parafilm to allow ethanol to absorb and then seeded with OP50 bacteria. NAC plates: NAC was added to NGM agar media after cooling to 55°C at a final concentration of 10 mM; plates were dried before seeding with OP50. 3HB plates: Ethyl-3-Hydroxybuterate was added to NGM agar media after cooling to 55°C at a final concentration of 50 mM; plates were dried before seeding with OP50. AA plates: Ethyl-Acetoacetate was added to NGM agar media after cooling to 55°C at a final concentration of 25 mM; plates were dried before seeding with OP50. FA plates: Folic acid (final concentration10 uM) and sodium bicarbonate (final concentration 5mM) were added to NGM agar media after cooling to 55°C; plates were dried protected from light and seeded with OP50 once dry. Control plates with NGM with 5mM sodium bicarbonate were used as controls for FA plates.

### Hatch assays

To determine rates of embryonic viability, two L4 hermaphrodite worms were plated on a 60 mm plate. Sixteen to twenty-four hours later, the now gravid adult hermaphrodites were removed from the plate and the number of eggs present on the plate were counted (any larvae present were ignored). After an additional 16-24 hours, the number of eggs present were counted again. Fraction hatched = (Day 1 count – Day 2 count)/Day 1 count. Empty eggshells are easily distinguished from arrested, or unhatched embryos.

### Imaging

Time lapse imaging data of fluorescent *C. elegans* embryos was collected with a Nikon A1R inverted LUNV confocal laser scanning microscope in the CCHMC Bio-imaging and Analysis Facility as previously reported^24^. Embryos that reached three-fold elongation and were moving after imaging were considered to have hatched. Widefield images of larva were collected with a Nikon Ni-E upright fluorescent microscope.

### Quantitative analysis

Hatch assays were analyzed with a two-sided binomial proportions test using R statistical software. Graphs were plotted using Prism software. For time-lapse confocal imaging of embryos, data were processed with StarryNite and manually curated with AceTree^35,36^. Curated nuclei positions and divisions were aligned and rotated using procedures previously outlined and cell displacement distance and nearest neighbor (NN) scores were calculated for each cell at each timepoint; the mean displacement and NN score were used for analysis and normalized to the average value each wildtype cell from the control dataset as some cells have intrinsically greater variability in position^23,24^. Cell position scores were calculated by adding the normalized mean displacement to the normalized mean NN score. Data for cell path length and cell cycle length were compared using a students t-test.

## ACKNOWLEDGMENTS

We thank Matt Kofron and the BAF staff for their assistance with imaging, John Murray for sharing software updates, and Marian Walhout and her laboratory staff for assistance with plate recipes. We thank the members of the Penn Worm Group and Brian Gebelein for support and constructive feedback. We thank Jichao Chen and Jeff Whitsett for financial support and acknowledge funding from a CCHMC Trustee Award and NIH R00 GM111825 to ALZ.

## FIGURES

**Supplemental Figure S1.**
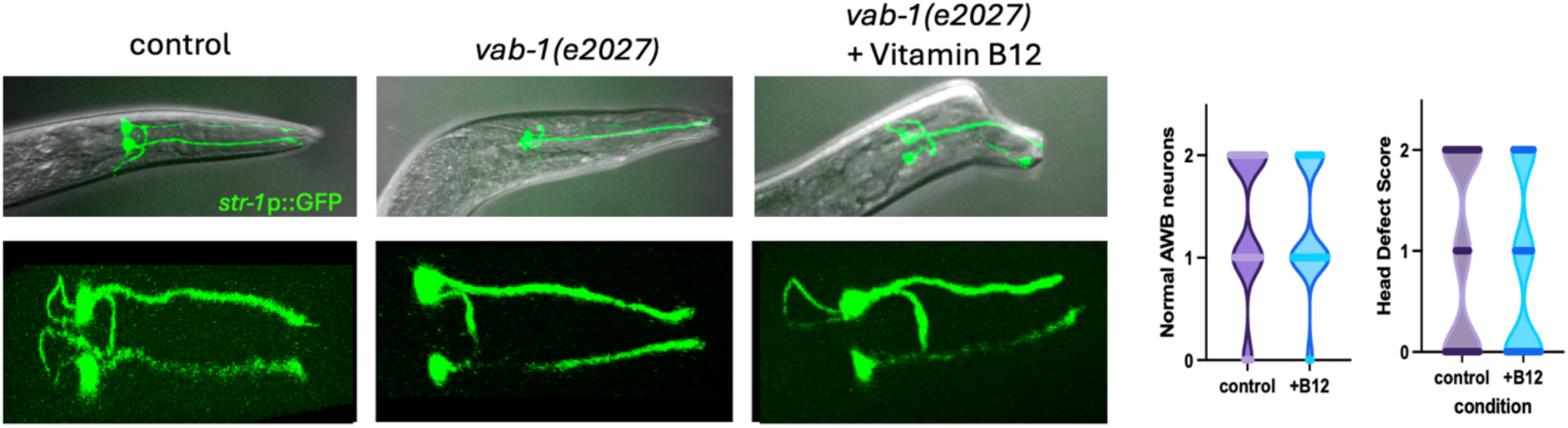
Vitamin B12 does not rescue larval defects in axon pathfinding or head morphogenesis. Widefield microscopy of L4 larva shows that *vab-1* mutant embryos frequently show misshapen heads due to disruptions in epithelial morphogenesis and loss of loops on the AWB neurons highlighted by the str-1p::GFP reporter. Quantification shows that the number of normal AWB neurons and severity of head defects (0=normal, 1=moderate, 2=severe) do not change with supplementation of vitamin B12. Worms were raised on B12 treated plates after hatching from a hermaphrodite parent which was transferred to the B12 treated plate at L4 stage.

**Supplemental Figure S4.**
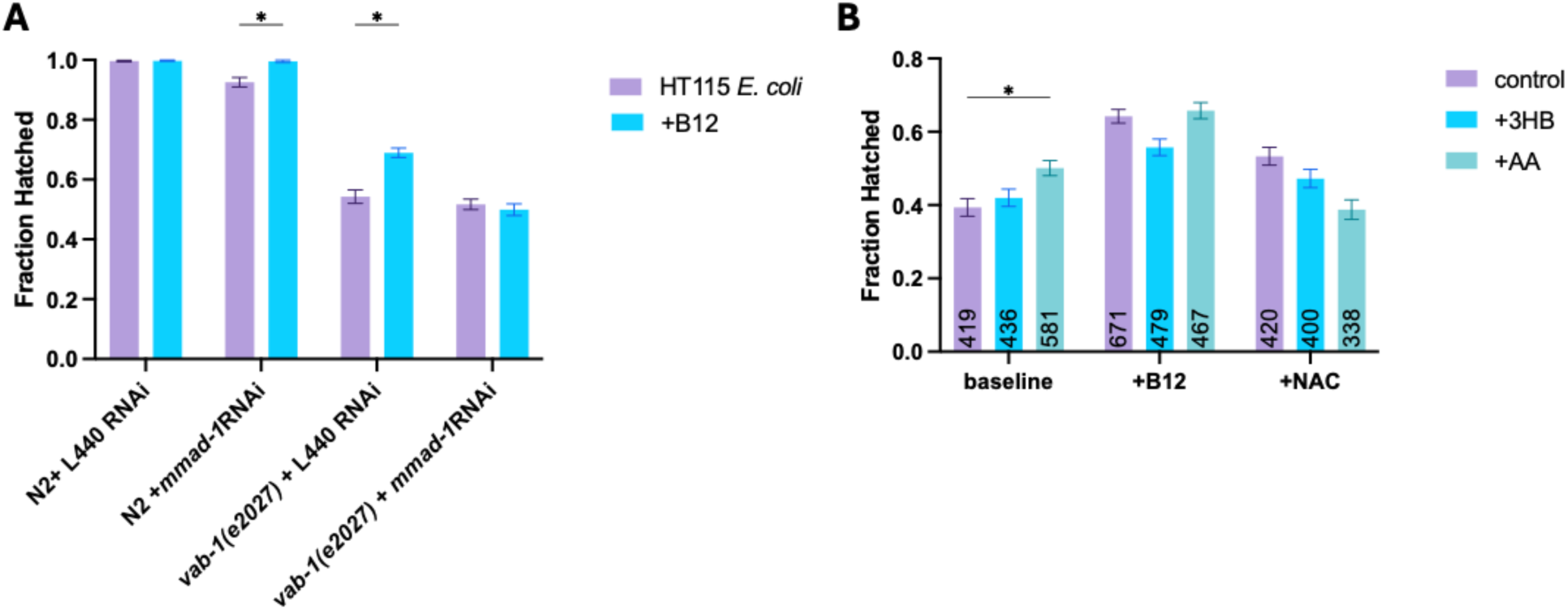
Vitamin B12 rescue depends on *mmad-1* function and cannot be effectively replaced by ketone body supplementation. A) Loss of the vitamin B12 processing enzyme *mmad-1* prevents rescue of *vab-1* embryonic lethality by vitamin B12. L440 empty vector in the HT115 *E*.*coli* strain was used as a control food source. B) Impact of ketone body supplementation on *vab-1(e2027)* embryo hatch rates when added to standard NGM plates, or those supplemented with vitamin B12 or NAC.

